# Neutrophils from Alzheimer’s Disease mice fail to phagocytose debris and show altered release of immune modulators with age

**DOI:** 10.1101/2025.04.21.649766

**Authors:** Matthew Kays, Anna Kelly, Farrah McGinnis, Clara Woods, Candice Brown, Aminata P Coulibaly

## Abstract

Recent reports show that neutrophil activity plays a role in the cognitive decline associated with Alzheimer’s Disease (AD). There is evidence of altered functions in neutrophils isolated from AD patients. Whether these altered functions are inherent to the AD disease state is unknown. The goal of this study was to determine if neutrophil functions are altered in AD mice and if these changes occur only after symptoms appear. To address this hypothesis, we used a primary neuronal culture model, generated from 3xTg perinatal mice, since AD is considered a CNS disease. The 3xTg primary neuronal culture gradually increase the release of Aβ (40 and 42) as the culture ages. To assess neutrophil functions, neutrophils isolated from young male/female mice (3-6 months of age) or aged male/female mice (16-18 months of age) of WT or 3xTg mice were exposed to 3xTg primary neuronal cultures. To assess phagocytosis, we characterized the effect of neutrophils on pathogenic amyloid beta (Aβ) 42 levels. To assess the levels of immune modulators (cytokines, chemokines, growth factors, NETosis, and neutrophil granular content), culture media were assessed using Luminex multiplex assay. Our results show that neutrophils from young AD mice have impaired phagocytosis, as observed in a decreased ability to remove Aβ and cellular debris in vitro. Neutrophils from young AD mice also increased release of pro-inflammatory cytokines, granule content, and NETs in 3xTg primary neuronal cultures. Interestingly, neutrophils from aged 3xTg mice decreased Aβ levels in culture and expression of proinflammatory cytokines when compared to neutrophils from aged WT mice. These neutrophils increased their release of granule content and NETs in 3xTg primary neuronal cultures. These data show that in AD neutrophil function is altered both prodromal (young mice) and diseased (old mice) stage.

## Introduction

With the increasing incidence of Alzheimer’s Disease (AD), there is dire need to fully understand its etiology with the hope of finding effective treatments. It is understood that AD is a progressive disease with onset long before cognitive symptoms manifest (prodromal stage). AD symptomology is characterized by abnormal cognitive and behavioral changes. These changes, including memory loss, diminishes a person’s quality of life and contribution to society. AD is characterized, pathologically, by the accumulation of two key proteins, extracellular amyloid beta (Aβ) and intracellular hyperphosphorylated Tau (p-Tau) (1). These two proteins have been shown to be highly toxic (2). There is mounting evidence of peripheral immune recruitment to the brain in AD (3). Neutrophils, a peripheral immune cell type, are found around Aβ deposits in the brain of AD patients (4). Several recent studies have demonstrated neutrophil recruitment to the brain vasculature in several rodent AD models (5–7). Interestingly, two studies have shown that neutrophils isolated from AD patients show impaired phagocytosis and increased release of inflammatory molecules (8,9). These data suggest that in AD, neutrophil functions are altered. Whether these cellular changes are present in pre-clinical AD models or occur in the prodromal phase of the disease are currently unknown. In this study, we hypothesized that neutrophil from Alzheimer’s Disease pre-clinical models show impaired function and activity in the prodromal and disease stages.

To test this hypothesis, we used the transgenic 3xTg mouse model of AD. This model contains 3 mutations associated with familial forms of AD. These mutations are the Swedish mutation in the amyloid precursor protein (K670N/M671L), a mutation in the presenilin protein (M146V), and a mutation in the Tau protein (P301L). The 3xTg mouse is slow progressing with Aβ detection, cognitive decline, and neuroinflammation at 6 months of age(10). Using cell culture, histology, cytokine assays, and amyloid ELISA, our data showed that neutrophils from 3xTg mice failed to phagocytose Aβ and cellular debris, showed altered cytokine production, and differed from WT derived neutrophils in both young (prodromal) and aged (diseased) mice.

## Material and methods

### Animals

Young (3-6 months) and aged (16-18 months), male and female 3xTg (B6;129-Tg(APPSwe,tauP301L)1Lfa *Psen1*^*tm1Mpm*^/Mmjax), C57Bl/6 wildtype, and CVN (APPSwDI-Nos2^-/-^) mice were used for this project. 3xTg and CVN perinatal day 0 were used for primary neuron culture generation. Mice were kept on a 12 hour:12-hour light cycle at room temperature (22- 25^°^C). Food and water were provided *ad libitum*. All experiments were done with the approval of West Virginia University Animal Care and Use Committee.

### Primary Neuron cultures

Brains of 3xTg and CVN P0 pups were collected to generate primary cortical neuronal cultures as previously described (11). Brains were removed from the pups. Brainstem and midbrain structures were discarded. Remaining brain sample was digested in a medium containing DMEM/F12, glucose, Glutamax, Na+/pyruvate, FBS and penicillin/streptomycin. Brain samples were centrifuged and resuspended in the same media. 12 well plates, coated with poly-D-Lysine, were seeded at a density of 2×10^5^ cells/well. After a 2-hour incubation, media was removed and replaced with media containing Neurobasal, Glutamax, B27, and penicillin/streptomycin. Cultures were maintained for 4 weeks with one third media replacement every 3-4 days. Starting at week 2, 500μL of media was collected from weekly each well and frozen for Aβ ELISA and Luminex assay.

### Neutrophil isolation

Neutrophil isolations were conducted using bone marrow from 3xTg, CVN, and wildtype mice at timepoints specified above. The animals were sacrificed using isoflurane and bone marrow was obtained from the tibia and femur. Neutrophils were isolated using the MojoSort Neutrophil Isolation Kit (BioLegend, 480058), according to the manufacturer’s protocol. The MojoSort kit enriches neutrophils to 80% in our bone marrow isolate (Sup Fig 1A). Isolated neutrophils were added to the primary neuron culture at the end of week 4. Neutrophils were seeded at a density of 2×10^5^ cells/well. Co-cultures were incubated for 6 hours. Media was collected from each well after the incubation period, and plates were fixed with 4%PFA for IHC staining.

### ELISA assays

ELISA plates for Aβ40 (Abcam, ab193692) and Aβ42 (ThermoFisher, KHB3441) were used to assess soluble Aβ levels in supernatant collected from each pup throughout the four weeks. The manufacturer protocols for the Aβ42 ELISA and Aβ40 ELISA kits were followed. Briefly, antibody solutions were added to all wells (standards, controls, and samples) and incubated for 3 hours. The wells were washed, HRP-Streptavidin solution added, and incubated for 45 minutes. The plates were then washed, and stabilization chromogen added. Plates were read using a plate reader (SpectraMax iD3) at 450nm absorbance.

### Luminex

15-plex Luminex assays (bio-techne, CUSTOM-LXSA-M15) were used to measure secretions of immune modulators in supernatants collected from co-cultures experiments. The proteins assessed were pro-inflammatory cytokines (IL-1beta, IL-6, IL-17, TNFalpha, IL-alpha), anti-inflammatory cytokines (IL-4, IL-10), chemokines (CCL2, CXCL1, CXCL12), growth factors (G-CSF, IGF-1, beta-NGF), a marker for degranulation (MMP-9), and a marker for NETosis (S100A8). Luminex assays were run according to manufacturer’s protocol by the Flow Cytometry and Single Cell core facility at West Virginia University Health Science Center.

Assays were run on the Luminex MAG PiX instrument using the Xponent software version 4.3.

### Free DNA detection

The concentration of free DNA in supernatants from the cultures was determined using the Quant-iT PicoGreen dsDNA assay kit (ThermoFisher, P11496). Assay was performed according to manufacturer’s protocol. Plates were read using a plate reader (SpectraMax iD3) set to fluorescence at 480 excitation and 520 emissions.

### Immunohistochemistry Staining

Paraformaldehyde fixed culture plates were stained using basic immunofluorescence staining protocols. Briefly, cultures were incubated in blocking buffer containing 5% normal goat serum and 0.1% Triton X in 1xPBS for 30 minutes; followed by incubations with primary antibody cocktails (Aβ37-42 (Cell Signaling, 8243), neutrophils (Ly6B; ThermoFisher, MA5-16541), neurons (beta III Tubulin; Abcam, ab78078), and astrocytes (GFAP; ThermoFisher, PA1-10004)) for two hours. Cultures were subsequently incubated with the appropriate secondary antibody cocktail (ThermoFisher) in 1xPBS one hour. Cultures were imaged on the Echo microscope, converted to multi-channel images using ImageJ, and quantified using IMARIS.

### Quantification using IMARIS

Images were analyzed and quantified using IMARIS. 3D reconstructions were made using the surface analysis tool on IMARIS. 3D reconstructions were used to determine the number of neutrophils (Ly6B+) and the size distribution (fragment area, µm^2^) of amyloid beta staining. Internalized and non-internalized Aβ fragments were determined by the distance between the neutrophil surface and the Aβ surface. Surfaces with distances more than 1µm away from neutrophil (Ly6B+) edge were considered outside of neutrophils and amyloid beta surfaces below 0 µm (0 indicated the edge of the neutrophil surface) were labeled inside neutrophils.

### Statistical Analysis

All statistical analyses were performed using GraphPad Prism. One-way ANOVA was used for comparisons among the control and neutrophil groups for ELISA data. A two-way ANOVA was used for comparisons among the control and neutrophil groups for the size distribution of internalized Aβ. A one-way ANOVA with repeated measures was performed on raw concentration values for each analyte from the Luminex assay. Significance was ascribed at p<0.05.

## Results

### Primary neuron cultures from 3xTg mice release increasing levels of Aβ in vitro

Previous studies showed very little astrocyte contamination of primary neuronal cultures generated from P0 pups(12), with cortical cultures having 4% and hippocampal cultures 28% astrocytes (GFAP) contamination(12). Therefore, we characterized the GFAP content observed in our cultures. In cultures generated from 3xTg pups, GFAP+ cells made up on average 15% of cells in cultures (Sup Fig 1C). To determine if 3xTg primary neuronal cultures release Aβ in-vitro, Culture media were collected at 2, 3, and 4 weeks in vitro. Our data showed that as early as 2 weeks in vitro, primary 3xTg neuronal cultures release detectable levels of Aβ (Fig 1A). Characterization of two Aβ species, show a gradual increase of Aβ-42 (one-way ANOVA: p=0.013, F(1.06, 17.01)=7.31) and steady levels of Aβ-40 (Sup Fig 2A.). Aβ-42 levels were significantly increased at both 3wks (p=0.01) and 4wks (p=0.02) from the 2wks time point in-vitro (Fig 1A).

**Figure 1:**
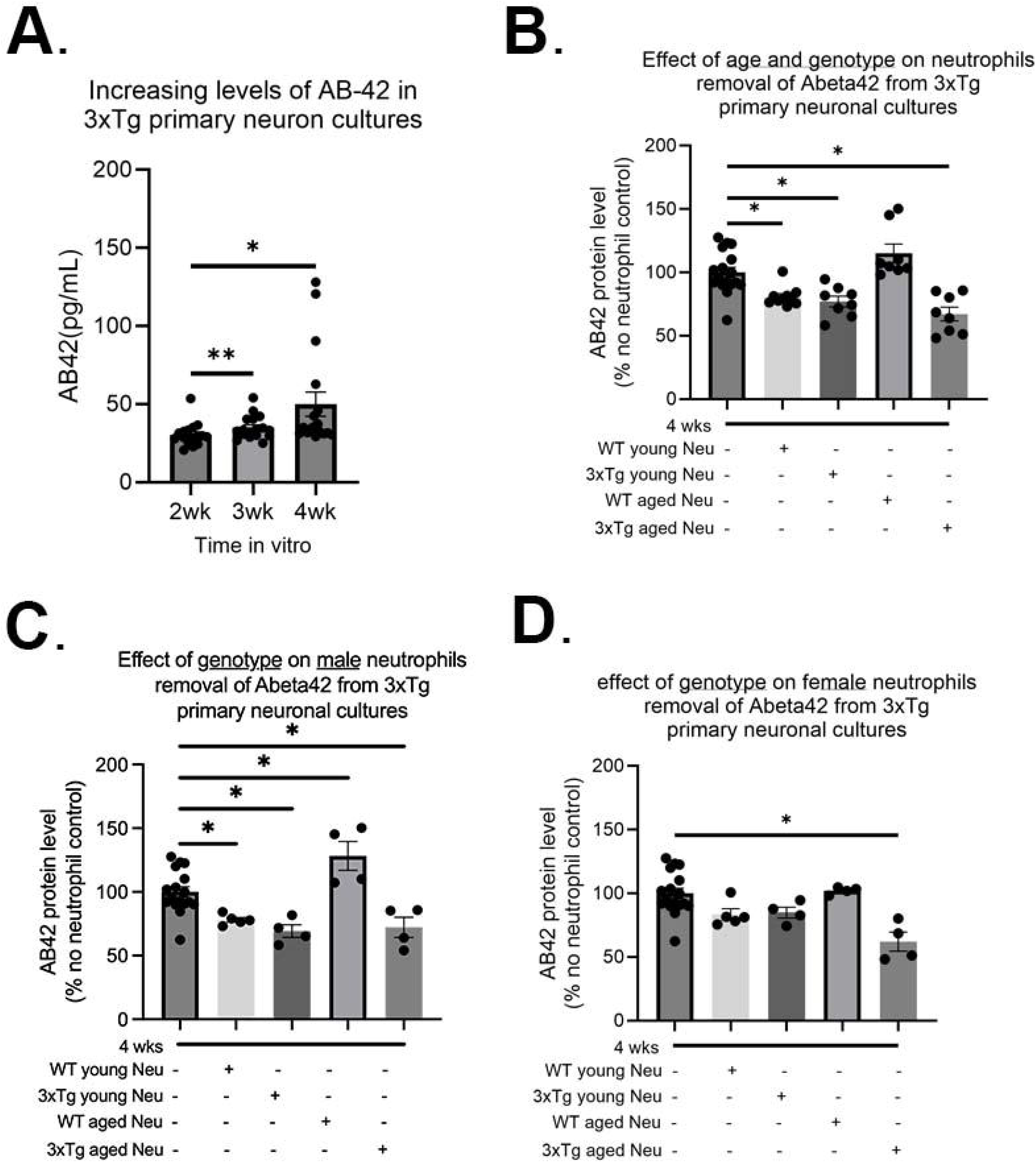
Both genotypes and sex affect soluble Aβ-42 removal from 3xTg primary neuronal cultures. Primary neuronal cultures, generated from P0 3xTg mice, produced increasing levels of Aβ-42 over 4 weeks in vitro (A.). The addition of bone marrow neutrophils from young and aged mice significantly decreased Aβ-42 levels in 4week cultures, except those derived from WT aged mice (B.). Male neutrophils from WT/3xTg young and 3xTg aged neutrophils decreased the level of Aβ-42 while WT aged neutrophils increase its levels (C.). For female mice, only neutrophils from aged 3xTg mice significantly decreased the levels of Aβ in cultures. Each dot represents data from one pup. *, p<0.05.

### Neutrophils from female 3xTg mice fail to reduce soluble Aβ42 levels in culture

Of note, the bone marrow of 3xTg mice contain significantly less neutrophils than WT bone marrow (Sup Fig 1D). Specifically, the number of neutrophils in young 3xTg bone marrow was similar to the number observed in aged WT mice (Sup Fig 1D). To determine whether the presence of neutrophils would affect the levels of Aβ in the 3xTg primary neuronal cultures, at 4 wks, in vitro cultures were exposed to neutrophils from WT and 3xTg mice. The addition of neutrophils significantly decreased Aβ-42 levels (One-way ANOVA: p <0.0001; F(4,46)= 14.29; Fig 1B) with no change to Aβ40 levels (sup Fig 1B). Further characterization showed that neutrophils from young WT (p=0.010), young 3xTg (p=0.003), and aged 3xTg (p=0.00002) significantly decrease Aβ-42 levels, while those from WT aged mice did not (Fig 1B).

To determine if sex or age influenced the effect of neutrophils on Aβ levels, cultures were incubated with neutrophils isolated from male or female mice from WT/3xTg, young/aged mice. For male derived neutrophils, cells from WT young (p= 0.039), 3xTg young (p= 0.005), and 3xTg aged (p=0.013) significantly decreased Aβ-42 levels in culture (Fig. 1C). Interestingly, neutrophils from WT aged male mice increased the levels of Aβ-42 levels in-vitro (p= 0.011) and those from 3xTg young male mice increased Aβ-40 levels (sup Fig 1C). On the other hand, only neutrophils from 3xTg aged female mice significantly decreased Aβ-42 levels (p=0.0001; Fig. 1D). These data suggest that the activity of neutrophils changes based on the disease state and sex of the mouse.

### Neutrophils from young 3xTg mice fail to phagocytose Aβ plaques and debris in culture

To assess the activity of neutrophils in the 3xTg primary neuronal cultures, we characterized phagocytosis by determining the percentage of neutrophils with internalized Aβ fragments, the size distribution of internalized fragments, and the number of Aβ plaques (Fig 2A). Most Aβ plaques were found outside of beta-3-tubulin+ neurons (Fig 2A). On average 60% of neutrophils from young mice contain Aβ fragments (Fig 2B&D), which was significantly increased to 75% in neutrophils from young 3xTg male mice (Fig 2B; p=0.0287). Neutrophils from young male 3xTg mice contained fewer Aβ fragments than those from WT male mice (Fig 2C; p=0.0213). To determine if this internalization led to a decrease in plaque burden, we assessed the distribution and range of remaining Aβ plaques in culture. Neutrophils from young WT mice had fewer Aβ fragments internalized but showed the largest reduction in Aβ plaque number. This suggests a continuous internalization and breakdown of Aβ by these cells. On the other hand, neutrophils from young male 3xTg male mice failed to decrease the number of Aβ plaques in cultures, even with an increased number of internalized fragments. This suggest that these cells can internalize Aβ but may fail to degrade it, leading to a cytoplasmic accumulation over time. Neutrophils from female 3xTg had fewer internalized Aβ fragments compared to those from young WT female mice (Fig 2E; p=0.0022). Interestingly, neither decreased the distribution or number of Aβ plaques in culture. These data suggest that sex and genotype contribute to neutrophil internalization of Aβ in vitro.

**Figure 2:**
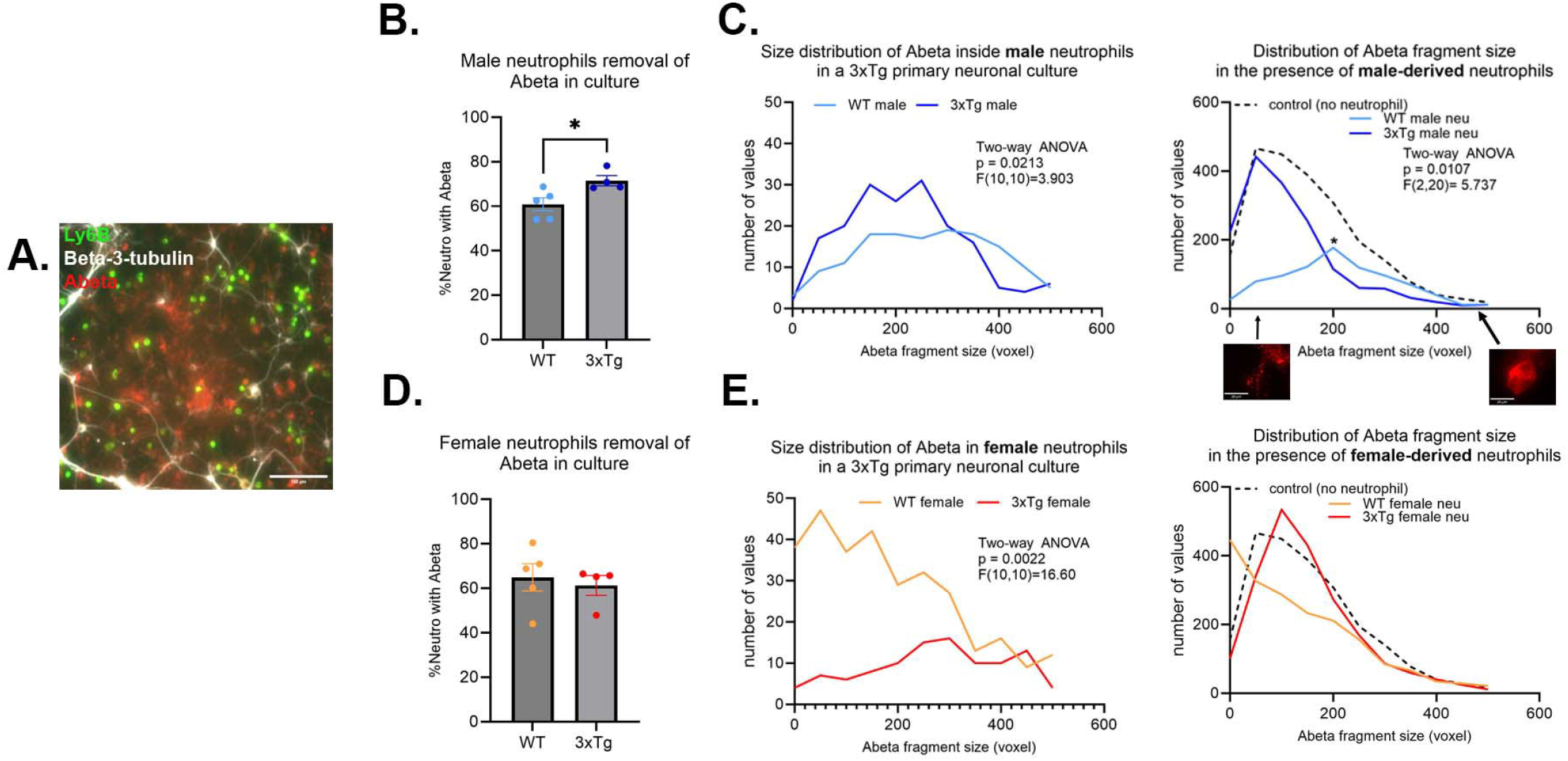
Neutrophils derived from young (3-6 months) WT mice are better at removing Aβ plaques in vitro. A representative micrograph depicting the presence of Aβ plaques (red) in 3xTg primary neuronal cultures (white; beta-3-tubulin) after the addition of isolated neutrophils (green; Ly6B) (A.). More neutrophils from young 3xTg male mice contained Aβ than those from young WT male mice (B.). Further analysis of the size of the Aβ fragments inside the cells show more fragments in the 3xTg derived neutrophils (C.). Overall, only young WT male neutrophils significantly decreased the number and Aβ plaques in vitro (C.). The number of neutrophils from young WT and 3xTg female mice with internalized Aβ was not different (D.). WT female neutrophils internalize more Aβ fragments than those from the 3xTg with no effect on the distribution and number of Aβ plaques in culture (E.). Scale bars: A= 100 µm; C=20 µm. Each dot represents data from a pup. *, p<0.05.

To determine if the deficit in phagocytosis is specific to Aβ, we assessed neutrophil removal of cellular debris. For this experiment, we generated primary neuronal cultures from the CVN-AD (13) mouse, a vascular model of AD(14). Primary neuronal cultures from this strain die at 3 weeks in culture, leading to a plate with neuronal debris and fragments. Similar to neutrophils from 3xTg young male mice, neutrophils isolated from the young male CVN mice fail to clear cellular debris (Sup Fig3). Neutrophils from young WT male mice remove a significant number of cellular debris in comparison (Sup Fig2; p<0.001). Neutrophils from young WT female mice removed cellular debris efficiently, while those from the young female CVN mice fail to do so (Sup Fig3; p<0.0001). These data further support an inherent deficit in function of neutrophils from AD mouse models.

### Neutrophils from aged mice fail to clear Aβ in a 3xTg primary neuronal culture

In young mice, ∼60-70% of neutrophils contained internalized Aβ fragments. This was significantly decreased in aged mice, with only ∼40-50% contained Aβ fragments (Sup. Fig4; Two-way ANOVA: p=0.0019, F (1,30) =11.63)). This suggests that age influences neutrophil phagocytosis regardless of disease state.

Characterization of neutrophils from aged mice showed an abrogation of the changes observed in the cells derived from young mice (Fig2). Specifically, the 3xTg specific changes observed in the male mice were gone (Fig 3A & 3B). On the other hand, neutrophils from 3xTg female mice, which showed no changes in the young mice, show a significant increase in the number of cells internalizing Aβ compared to neutrophils from aged WT mice (Fig 3C; p=0.0172). Interestingly, neutrophils from WT aged female mice led to an increase in Aβ plaques deposit (Fig 3D.; compared to no neutrophil control p=0.0010). These data suggest that age directly affects neutrophil ability to regulate Aβ levels.

**Figure 3:**
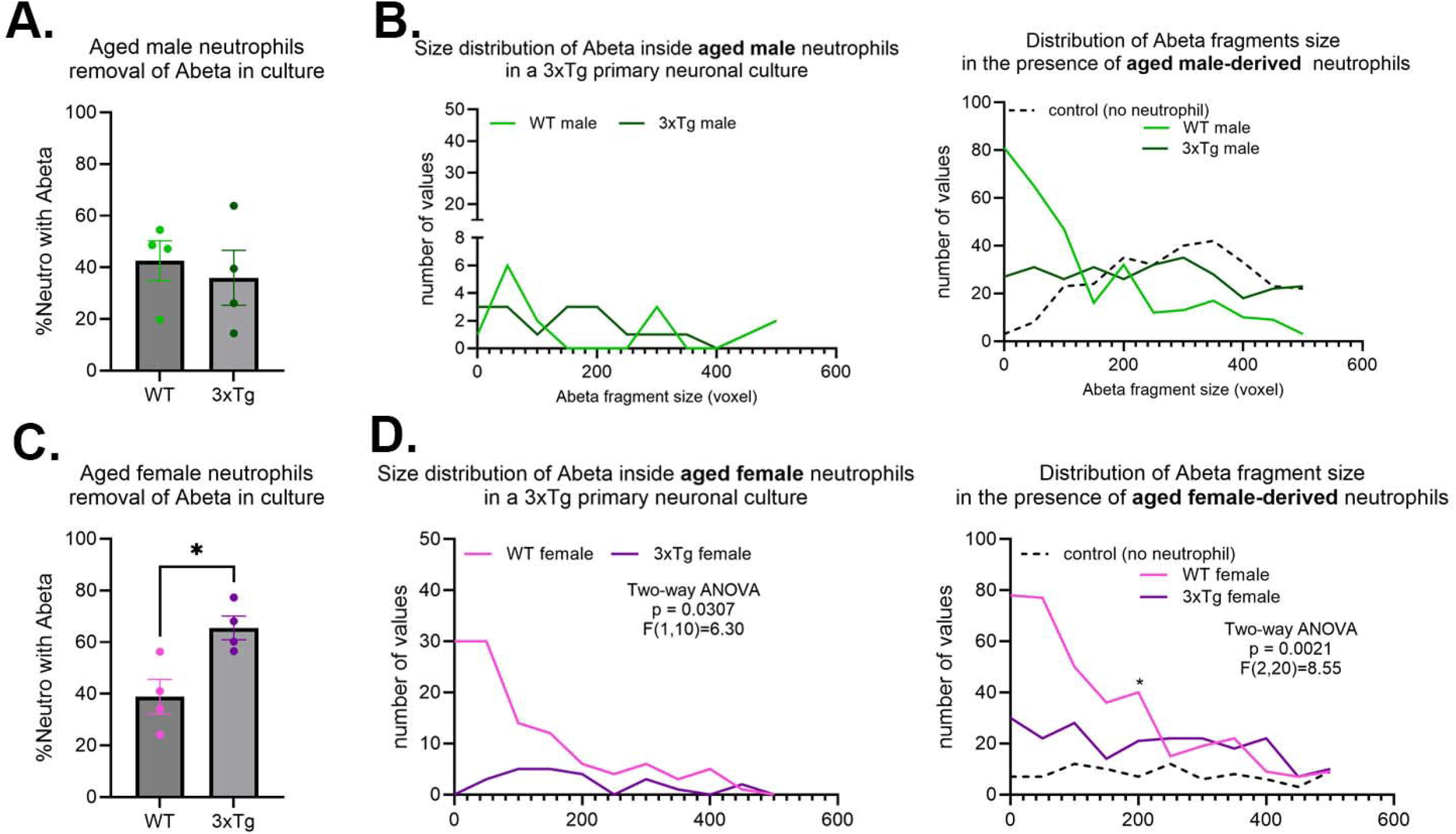
Age alters neutrophils ability to remove Aβ in vitro. No difference in neutrophils internalizing Aβ from WT or 3xTg aged mice (A.), with no change in the distribution and number of Aβ fragments inside aged neutrophils or their effect on overall Aβ plaques in cultures (B.). More neutrophils from aged 3xTg female mice contained Aβ fragments compared to those from the aged WT female (C.). Further analysis showed more Aβ fragments inside neutrophils from aged 3xTg female (D.). Interestingly, the presence of neutrophils from aged WT female mice increased the number of Aβ plaques in cultures (D.). Each dot represents data from a pup. *, p<0.05.

### Both age and animal disease state alter neutrophils release of immune modulators

Using a 15-plex immune array we characterized the production of pro-inflammatory cytokines, anti-inflammatory cytokines, growth factors, chemokines, and markers of degranulation and NETosis in 3xTg cultures exposed to neutrophils isolated from WT/3xTg, young/aged mice.

It is evident that 3xTg cultures exposed to neutrophils isolated from young 3xTg mice (male and female) have significantly higher levels of pro-inflammatory cytokines (IL1-b, IL1-a, IL6, TNF-a, IL17a) (Fig. 4A), and no change in the levels of anti-inflammatory cytokines (Fig. 4B). These cultures also showed increased presence of MMP9, a neutrophil granule protein (Fig. 4C), and increased NET production, as denoted by increased levels of S100A8 and free DNA (Fig. 4D). Cultures exposed to neutrophils from young WT mice had no effect on cytokines, MMP9, or NETs levels in the culture media. These data suggest that neutrophils isolated from 3xTg young mice generates an inflammatory environment in the presence of 3xTg neurons.

**Figure 4:**
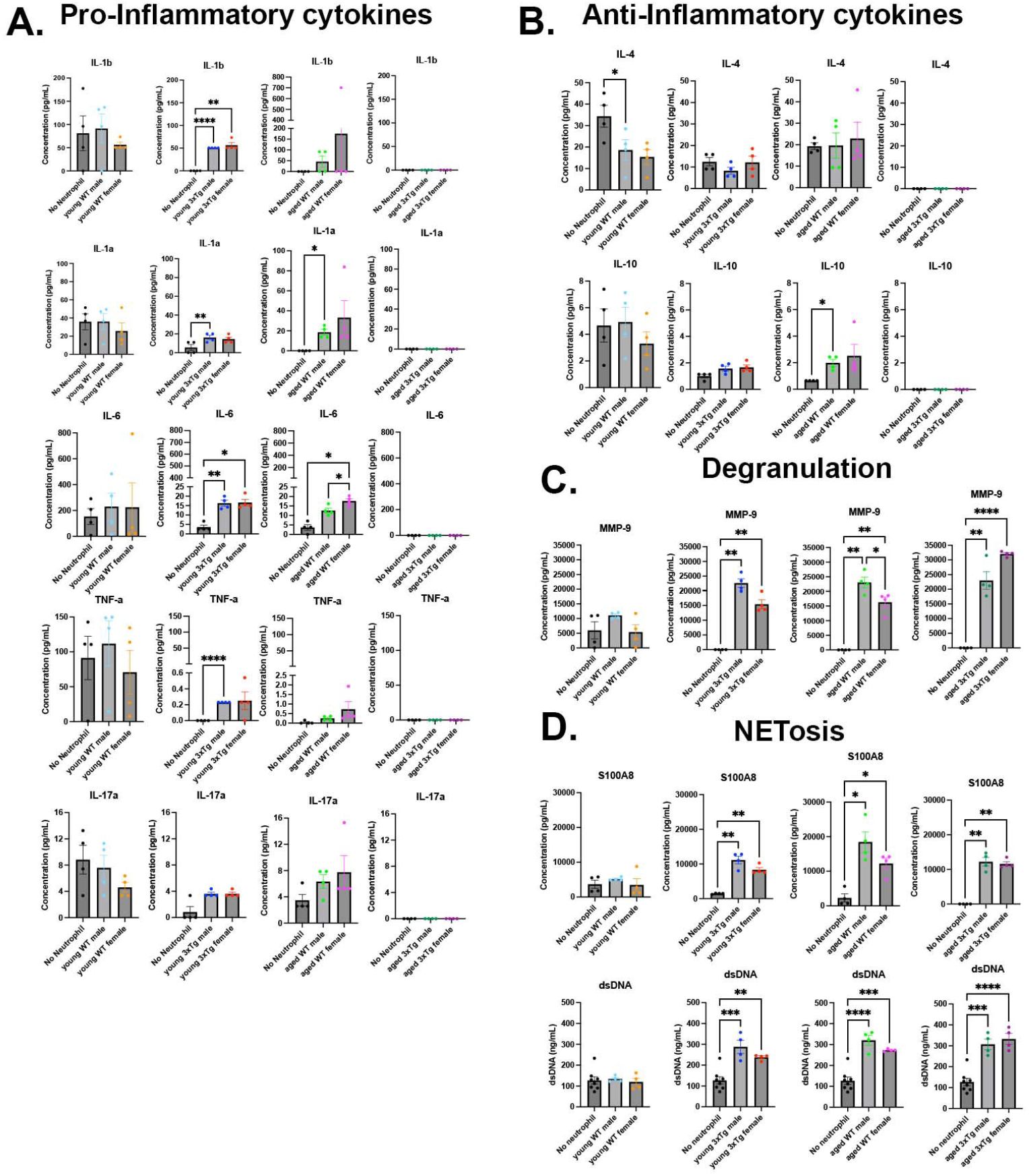
Neutrophils from 3xTg mice have altered release of immune modulators in 3xTg primary neuronal cultures. Culture media contained higher levels of pro-inflammatory cytokines with neutrophils isolated from young 3xTg mice and aged WT mice (A.). This increase is absent in cultures with neutrophils from young WT mice and aged 3xtg mice (A.). IL-4 is decreased when young WT male neutrophils are added to cultures while IL-10 is increased in the presence of neutrophils from aged WT male mice (B.). Markers of degranulation (C.) and NETosis (D.) were increased in cultures with all neutrophils except those from young WT mice. Each dot represents data from a pup. *, p<0.05.

On the other hand, exposure of 3xTg neuronal cultures to aged WT neutrophils led to an increase in pro-inflammatory cytokines, IL-6 and IL1-a, and the anti-inflammatory cytokine IL-10 (Fig. 4A&B). Interestingly, cultures with neutrophils from aged 3xTg mice showed no effect on cytokine levels, a direct opposite of what is observed in cultures with neutrophils from young 3xTg mice. Markers of degranulation, MMP-9, and NETosis, S100A8 and free DNA, were significantly increased in 3xTg cultures exposed to neutrophils from both aged WT and 3xTg mice (Fig. 4C&D). These data suggest that the 3xTg disease state significantly alters the activity and response of neutrophils in the presence of neurons with age.

Chemokines and growth factors were also detected in culture media. The levels of G-CSF increased in 3xTg cultures with the addition of young 3xTg male neutrophils (Sup Fig5A.). Young 3xTg male and female neutrophils both decreased the levels of CXCL1 while aged 3xTg female neutrophils decreased the level of CXCL12 in culture (sup Fig5B.).

## Discussion

The goals of the present study were to investigate the ability of neutrophils from AD mice to clear neuron derived Aβ and determine how age influences basic neutrophil function. Our data showed a clear sex difference in young neutrophils ability to interact with the pathogenic form of Aβ. Neutrophils from young male mice, regardless of disease state, remove more Aβ than neutrophils from young female mice. This sex difference is lost with age. In the aged-derived cells, only neutrophils from the 3xTg mice, male and female, significantly decrease Aβ levels. All 3xTg neutrophils showed impair phagocytosis regardless of age. Our data also show that neutrophils from young 3xTg mice release more pro-inflammatory cytokines, granule proteins, and NETs in 3xTg neuronal cultures than WT neutrophils. Neutrophils from aged 3xTg mice had increased release of NETs with no release of cytokines in 3xTg neuronal cultures, a result opposite to those from aged WT mice. These data clearly demonstrate that neutrophils from both young and aged AD mice have altered function from WT mice.

Neutrophils have been implicated in AD pathology. Neutrophils infiltrate the brain in various AD mouse models including APP/PS1, 3xTg, and 5xFAD (4,15,16). This infiltration was linked to increased microglia activity via release of CAP37 in 3xTg mice at 12 months of age (4). Our data show that neutrophils from aged 3xTg mice showed increased release of MMP9 and NETs, which led to increased microglia activity and inflammation (17). Neutrophil proteins co-localized with amyloid beta plaques (15). Neutrophils migrated towards amyloid beta plaques after infiltrating the AD brain (15). Our study showed that neutrophils not only interact but also phagocytose amyloid beta fragments. To our knowledge, we are the first to show, in vitro, that neutrophils can clear amyloid beta. Similar to our results, neutrophils from AD patients showed impaired phagocytosis of *staphylococcus aureus* or *Escherichia coli* (8,9). This suggest that AD disease state, in human and mice, dysregulates neutrophil physiology which may account for their aberrant activity in the disease. The underlying mechanism of this dysregulation is still unclear. Of note, our study showed that this defect is inherent to all AD neutrophils, not only those from the aged animal. Our data showed that aged 3xTg neutrophils decreased the concentration of Aβ without showing increased internalization. MMP-9 was shown to degrade Aβ peptides (18), suggesting that aged neutrophil release of MMP-9 could account for the decrease observed.

Neutrophils from aged AD patients (average age 67.4 years) and APP/PS1 mice (aged 12 months) show increased ROS production and NETosis (16) suggesting that neutrophils contributed to the inflammation observed in the AD brain. Though our data did not show an increase in cytokine production in neutrophils from aged 3xTg mice, a clear increase was observed in free DNA present in culture. Our data from young mice suggest a high inflammatory potential of these cells. It is interesting that this inflammatory profile diminished completely with age, whether this change affects disease progression is unknown. Finally, Although MPO was observed in plaques in human AD brains, neutrophil extracellular traps were only localized to the brain vasculature (16). Our data shows that in the presence of AD neurons, neutrophils from 3xTg mice release NETs. Since, neutrophils from WT mice do not release NETs in the presence of 3xTg neurons, we postulate that the increase NETosis could be an indication of further dysregulation of neutrophil function in 3xTg mice. Many reports of neutrophils in various AD model characterize these cells in the aged brain microvasculature (3,16). Therefore, an importance of the work presented here is that our data show that neutrophils from young AD mice, before symptoms occur, already show functional dysregulation. In addition, the dysregulation of bone marrow neutrophils, as shown in our data, suggests that AD, a brain specific disease, influences the activity of peripheral systems.

## Conclusion

This study showed that the neutrophils from AD mice are functionally different than neutrophils from WT mice. Our data showed a sex difference in how neutrophils interact with amyloid beta in the presence of neurons. The observed sex difference is age dependent (only present in neutrophils from young mice). Whether this contributes to the sex bias observed in AD (both mice and humans) remains to be determined. Our results showed that neutrophils from young and age 3xTg mice have altered interact with debris and release immune modulators. We postulate that neutrophil dysfunction underlies the detrimental contributions of neutrophils to AD pathology.

## Acknowledgements

The Luminex experiments listed here were conducted in the West Virginia University Health Sciences Center Flow Cytometry and Single Cell Core Facility (SCR_017738), which is funded by the TME CoBRE GM121322. Experiments that were relevant to the CVN mouse were funded by the NIH 1K22NS114363 awarded to Dr. Coulibaly.

---

**<u>Supplemental figure 1:</u> Validation of neutrophil isolation and 3xtg neuronal cultures**. The neutrophil isolation protocol enriched neutrophil purity to 80%(A.). Both beta-tubulin+ and GFAP+ cells were detected in primary 3xTg cultures (B.). GFAP+ cells were only15% of cells in culture on average (C.). More neutrophils were isolated from the bone marrow of young WT mice compared to young 3xTg mice and aged mice of both genotypes (D.).

**<u>Supplemental figure 2:</u> Neutrophil have little effect on levels of soluble A**β**-40 in 3xTg primary neuronal cultures**. 3xTg primary neuronal cultures showed little change in Aβ40 levels (A.). The addition of neutrophils to the cultures had no effect on Aβ-40 levels (B.). Only neutrophils derived from young male 3xTg mice increased the levels of Aβ-40 in culture (C.). Neutrophils from female mice had no effect on Aβ-40 levels (D.). Each dot represents data from a pup. *, p<0.05.

**<u>Supplemental figure 3:</u> Neutrophil from CVN mice, another AD rodent model, have impaired removal of cellular debris**. Neutrophils isolated from young CVN mice and incubated in culture plates with neuronal debris showed that neutrophils derived from male and female CVN mice fail to remove cellular debris when compared to their WT counterparts.

**<u>Supplemental figure 4:</u> Aging decreases the number of neutrophils phagocytosing A**β **in vitro**. Aged neutrophils showed a decrease in the percentage of neutrophils with internalized Aβ in 3xTg primary neuronal cultures. Each dot represents data from a pup.

**<u>Supplemental figure 5:</u> The presence of neutrophils had little effect on the levels of growth factors and chemokines in 3xTg primary neuronal cultures**. The addition of neutrophils from young 3xTg male mice increased the release of G-CSF in culture (A.) Young male and female 3xTg neutrophils decreased the release of CXCL1 in culture while aged 3xTg females decreased the release of CXCL12 (B.). Each dot represents data from a pup. *, p<0.05.

