## Supplementary figures and images for "Neutrophils from Alzheimer’s Disease mice fail to phagocytose debris and show altered release of immune modulators with age"

### sup fig 1

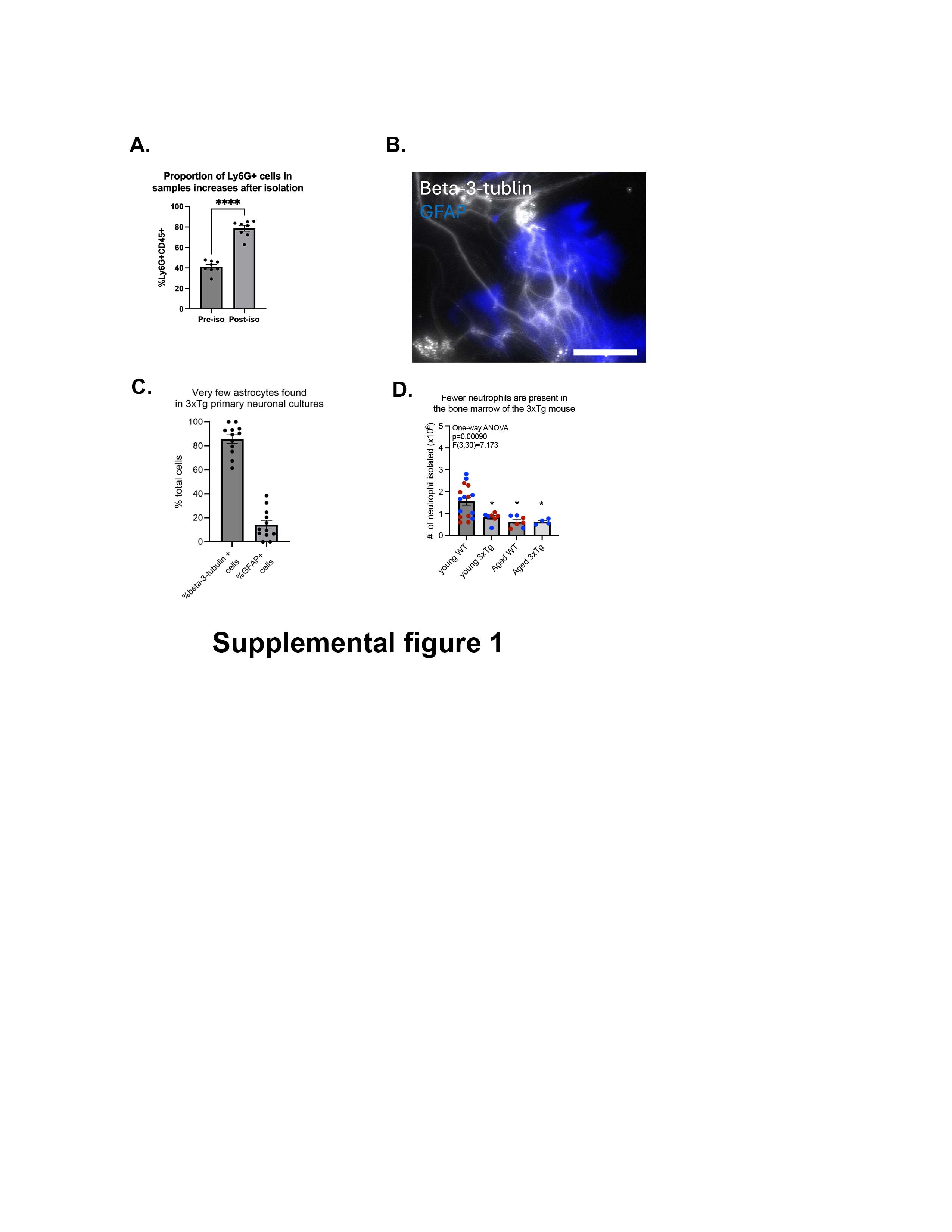

### sup fig 2

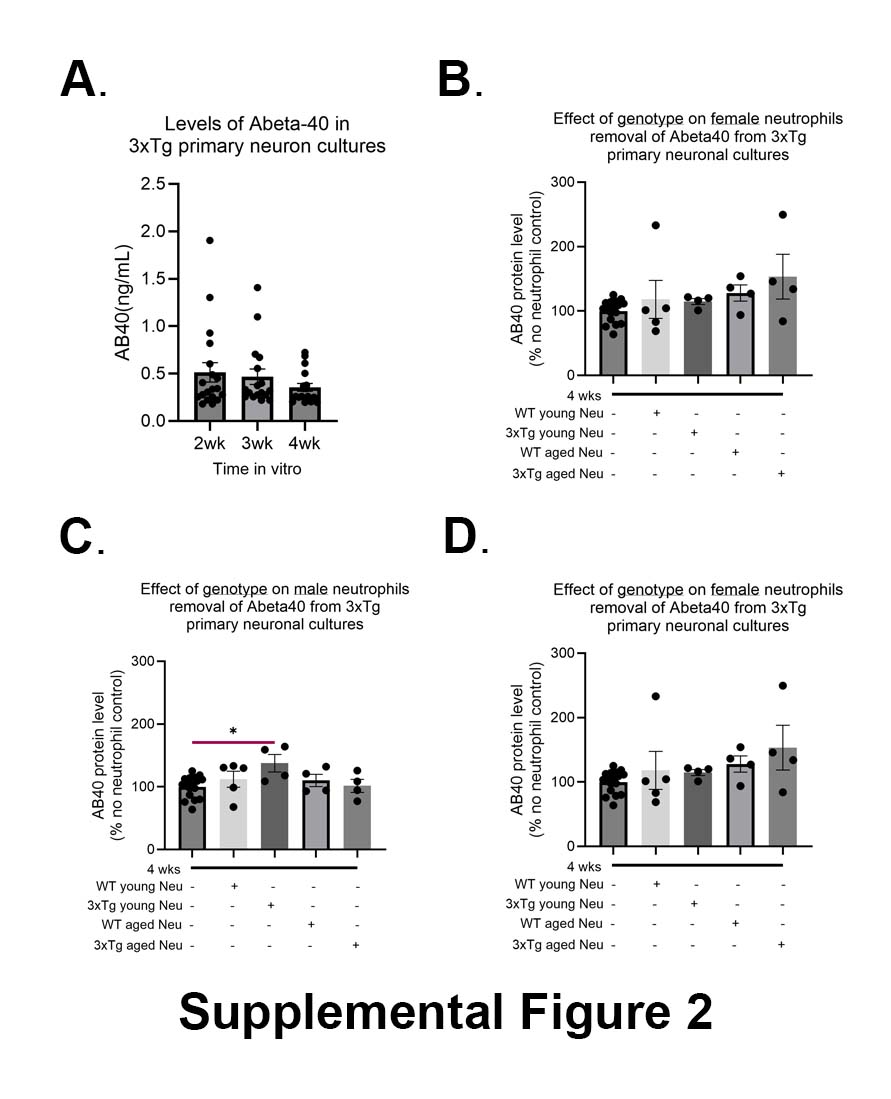

### sup fig 3

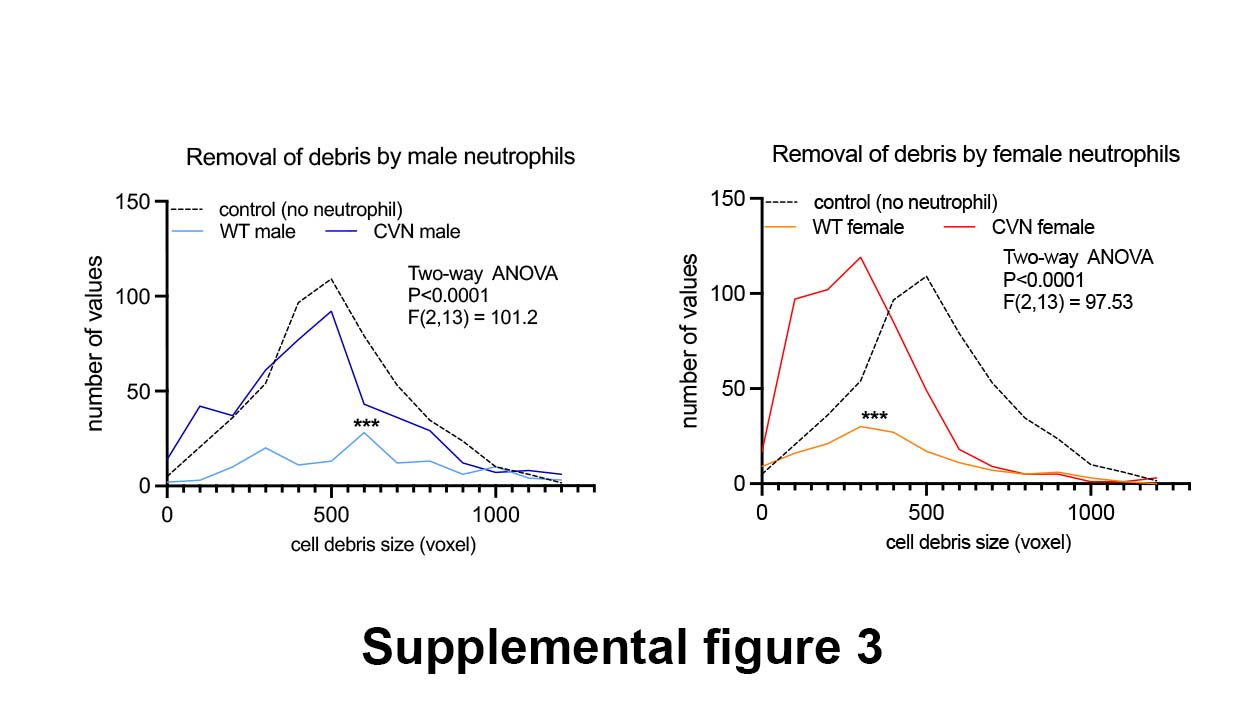

### sup fig 4

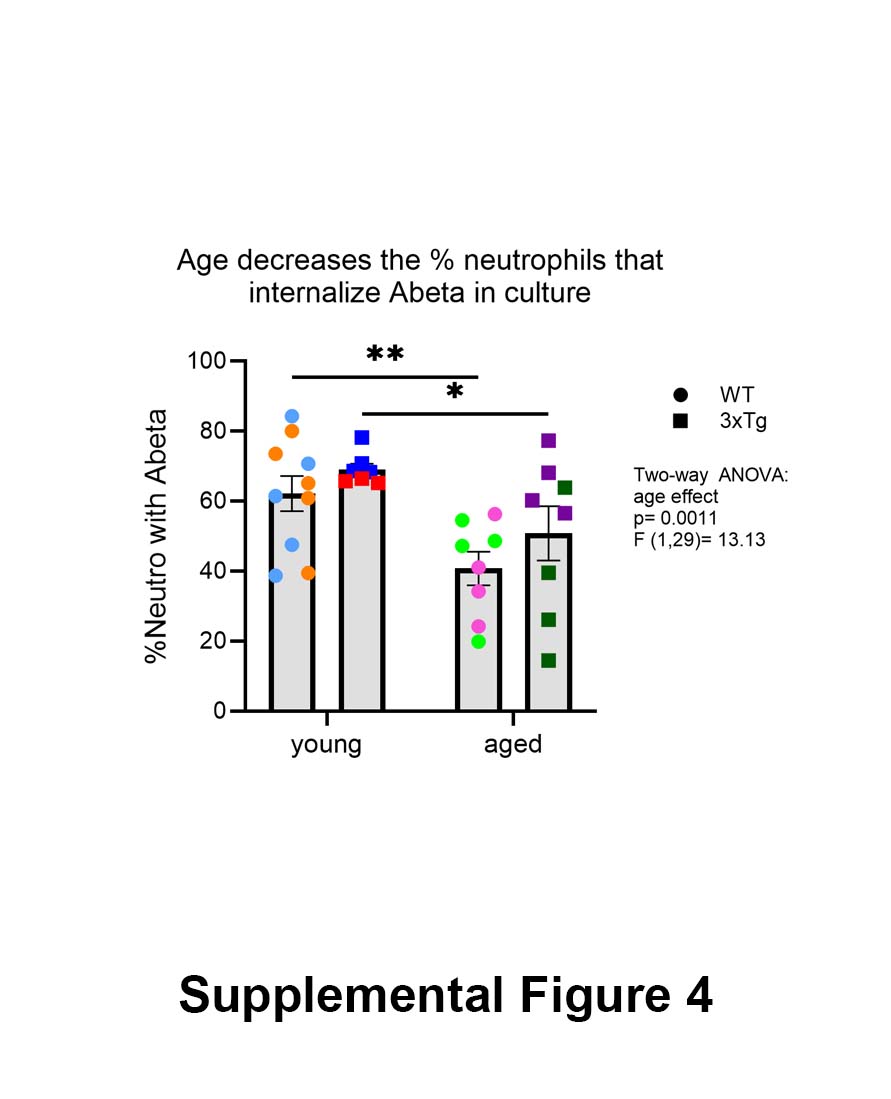

### sup fig 5

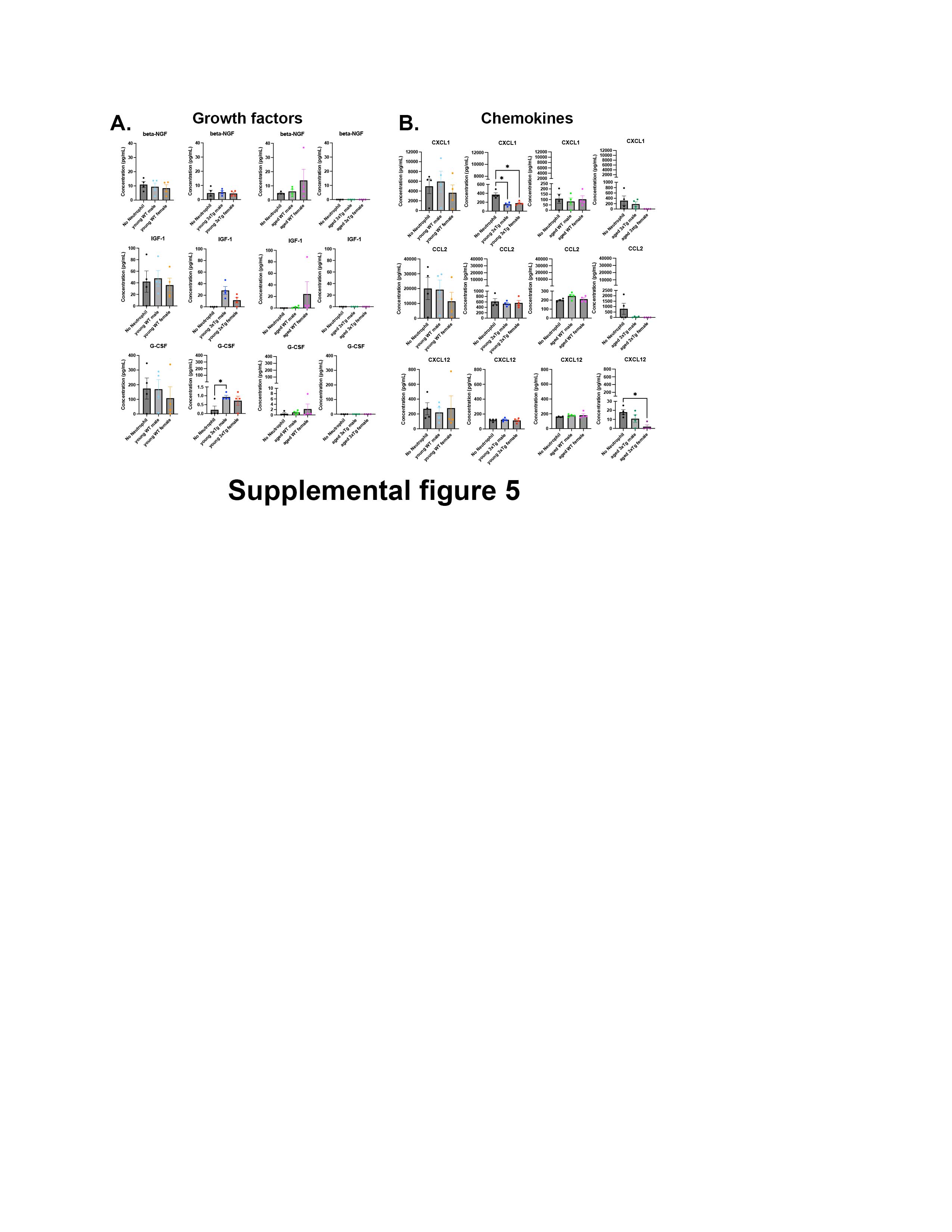
